# Defining epithelial stem cell heterogeneity through undulating structures of the skin and oral mucosa

**DOI:** 10.1101/2025.01.21.634195

**Authors:** Mizuho Ishikawa, Yen Xuan Ngo, Ikuto Nishikawa, Hiroko Kato, Ryo Maeda, Ryosuke Mizuno, Jun Mizuno, Kenji Izumi, Hiromi Yanagisawa, Aiko Sada

**Affiliations:** Division of Skin Regeneration and Aging, Medical Institute of Bioregulation, Kyushu University, Fukuoka, Japan; International Research Center for Medical Sciences (IRCMS), Kumamoto University, Kumamoto, Japan; Life Science Center for Survival Dynamics, Tsukuba Advanced Research Alliance (TARA), University of Tsukuba, Tsukuba, Japan; Ph.D. Program in Human Biology, School of Integrative and Global Majors, University of Tsukuba, Tsukuba, Japan; Graduate School of Medical Sciences, Kyushu University, Fukuoka, Japan; Graduate School of Medical Sciences, Kumamoto University, Kumamoto, Japan; Division of Biomimetics, School of Medical and Dental Sciences, Niigata University, Niigata, Japan; Research Center for Advanced Oral Science, School of Medical and Dental Sciences, Niigata University, Niigata, Japan; Taki Chemical Co., Ltd., Kakogawa, Hyogo, Japan; Komatsuseiki Kosakusho Co., Ltd., Suwa, Nagano, Japan; National Cheng Kung University, Tainan, Taiwan; Faculty of Medicine, University of Tsukuba, Tsukuba, Japan

**Keywords:** Epithelial stem cells, Oral epithelium, Skin, Proliferative heterogeneity, Rete ridge, Lineage tracing, Three-dimensional culture, Mechanical environment

## Abstract

Epithelial stem cells exhibit heterogeneity, with distinct stem cell populations occupying specific tissue regions. The human skin displays a characteristic undulating structure at the epidermal-dermal junction, which supports mechanical strength and influences the spatial organization of epithelial stem cells. Unlike human skin, mouse skin lacks these undulations, complicating studies into the effects of tissue architecture on stem cell distribution. Here, we leverage the mouse oral mucosa, which possesses an undulating structure similar to human skin, to characterize stem cell division dynamics and long-term fate *in vivo*. Using a combination of H2B-GFP pulse-chase analysis and lineage tracing with Dlx1-CreER and Slc1a3-CreER models, we demonstrate that slow-and fast-cycling stem cells localize to distinct anatomical regions relative to the undulating structure and maintain their respective compartments during tissue homeostasis. A three-dimensional culture model using micropatterned collagen scaffolds that recapitulate the undulating structures *in vitro* reveals that the mechanical environment generated by the undulating structures partially induces proliferative heterogeneity in epithelial stem cells. This study proposes tissue undulating surface structure as a common principle as a niche component that defines the localization of compartmentalized stem cell populations across different epithelial tissues.

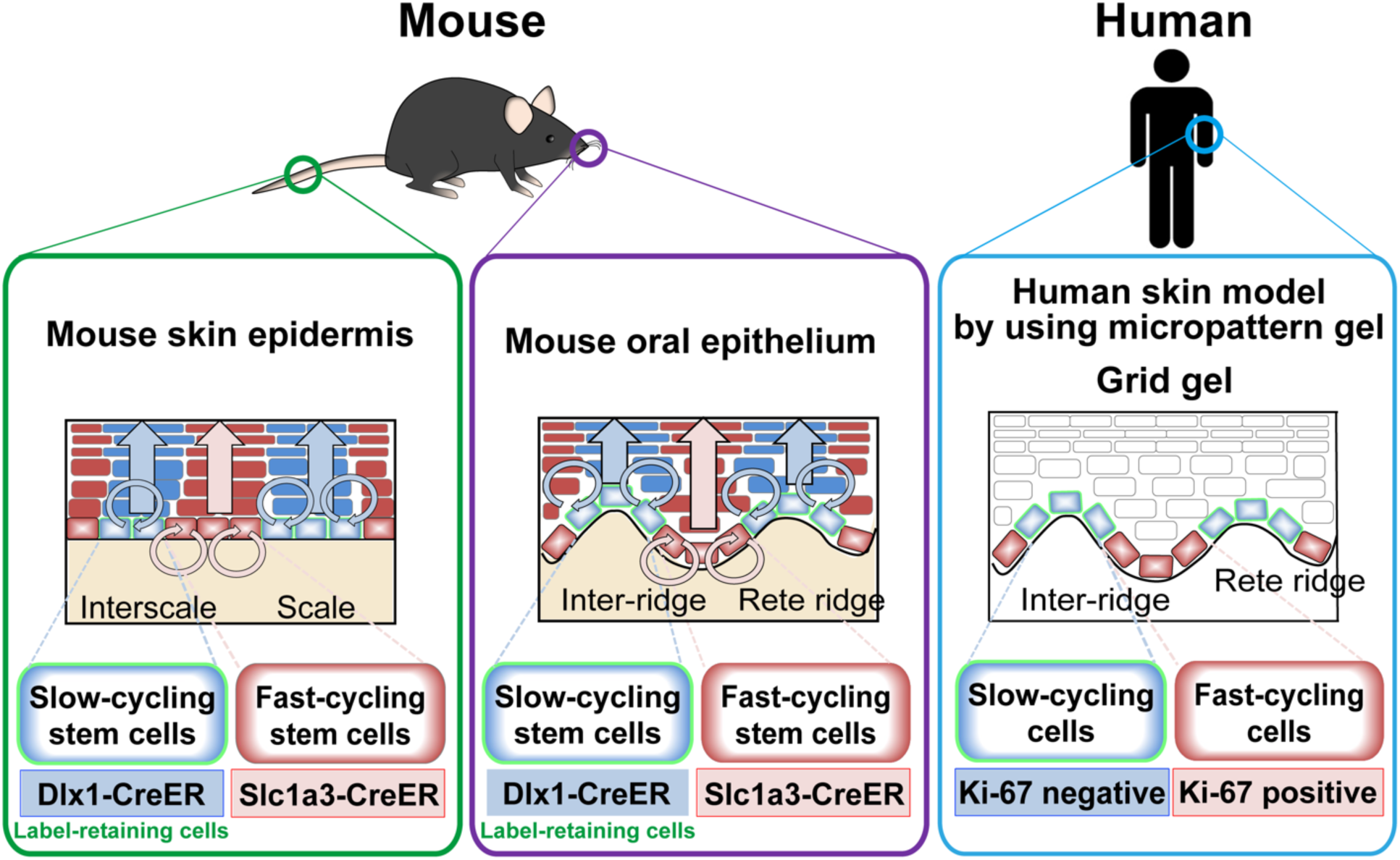

## Introduction

Epithelial tissue turnover and repair are sustained by tissue-resident stem cells. Studies have shown that epithelial stem cells are heterogeneous and localized to specific anatomical regions. In the intestinal epithelium, fast-cycling Lgr5+ stem cells are found at the crypt base, a region characterized by an undulating surface structure^1^. In the ocular surface epithelium, distinct slow-and fast-cycling stem cell populations occupy compartmentalized regions^2^. The limbus, a junction between the cornea and conjunctiva, enhances stem cell properties through mechanical forces associated with matrix stiffness^3^. Similarly, in the interfollicular epidermis of mouse tail skin, slow-and fast-cycling epidermal stem cells are localized in anatomically defined regions, namely the interscale and scale structures^4,5^. Recent single-cell RNA sequencing (RNA-seq) data from human skin suggests that slow-and fast-cycling epidermal stem cells are located in the rete ridges and inter-ridges (or dermal papillae), respectively, which form the undulating structures at the epidermal-dermal junction^6^. However, the role of tissue undulations in regulating epidermal stem cell identity and behavior remains unclear, as mouse skin lacks an analogous undulating structure for modeling.

The mouse oral mucosa features a rete ridge/inter-ridge structure, making it a unique *in vivo* model for studying tissue undulation. Oral epithelial cells expressing Bmi1 have been shown to divide rapidly and follow a population asymmetry model with neutral drift dynamics in the buccal mucosal epithelium^7^. In contrast, pulse-chase experiments have indicated proliferative heterogeneity in the gingiva, tongue, and palate^8–11^. The hard palate has an undulating rugae structure, which coincides with the heterogenous cell populations marked by Lrig1 and Igfbp5^12^. Recent spatial transcriptomics have revealed both unique and shared cellular niches and differentiation programs across distinct regions of the oral mucosa^13^. However, the relationship between the microscopic undulations of the rete ridge/inter-ridge structure and the heterogeneity of epithelial stem cells remains unclear.

An engineering approach has been employed to replicate the undulating structure of human skin *in vitro*. A polydimethylsiloxane (PDMS) scaffold with an undulating surface can partially induce the proliferation and differentiation patterns of epidermal stem cells in flat cultures of human keratinocytes^14,15^. Additionally, a patterned hydrogel has been shown to enhance stemness and activate signaling pathways in epidermal stem cells in a human three-dimensional skin model^16^. However, the influence of such undulating structures on epithelial stem cell heterogeneity remains unexplored.

In this study, we utilize mouse oral epithelium as an *in vivo* model to investigate the relationship between the undulating structure and the localization and behavior of epithelial stem cells. We also address how these structures affect stem cell heterogeneity by developing a three-dimensional culture system incorporating an undulating scaffold *in vitro*.

## Results

### Heterogeneous localization of slow-and fast-cycling cells along the undulating structure of the mouse oral epithelium

To investigate the relationship between the localization of epithelial cells with different division frequencies and the undulation structure, we performed an H2B-GFP label-retention assay in the mouse oral mucosa^17^. The oral mucosa consists of non-keratinized regions (ventral tongue, soft palate) and keratinized regions (hard palate; Fig. 1A, S1)^18,19^. In K5-tTA/pTRE-H2B-GFP mice, doxycycline treatment halts the transcription of histone H2B-GFP, leading to the dilution of the GFP signal with each cell division. This approach allows for the identification of slow-cycling cells as label-retaining cells (LRCs; Fig. 1B). At the 0-day chase, GFP labeling was uniformly present in oral epithelial cells (Fig. 1D-F). After a 2-week chase, LRCs were predominantly located at the inter-ridges, while non-LRCs were more frequently observed at the bottom of the rete ridges (Fig. 1G-I). Quantitative analysis confirmed that LRCs were enriched at distances 30 to 60 μm from the bottom of rete ridges, and that 20% of them were located in an area of more than 60 μm distance (Fig. 1C, J-L).

**Figure 1.**
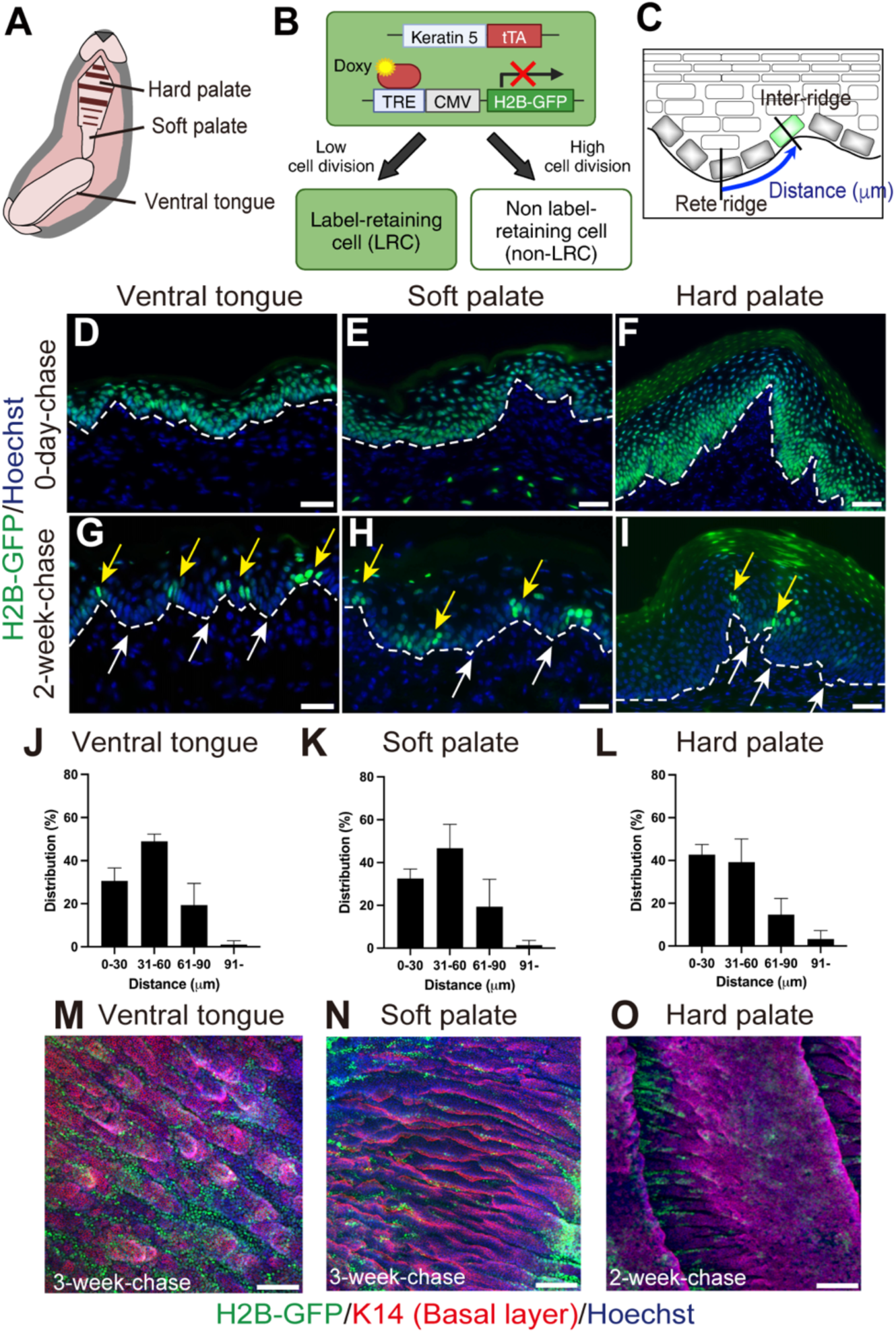
H2B-GFP label-retaining assay reveals the localization of slow-and fast-cycling cells in the mouse oral epithelium. **(A)** Schematic representation of the mouse oral mucosa. **(B)** Diagram of the K5-tTA tet-off/pTRE-H2B-GFP mouse model. Created with BioRender.com. **(C)** Quantification method for the distribution of H2B-GFP label-retaining cells (LRCs) in the oral mucosa. Distances from the base of the rete ridge to LRCs were measured. **(D-I)** Immunostaining of sagittal sections from the ventral tongue, soft palate, and hard palate at 0-day (D-F) and 2-week (G-I) chases. White arrows indicate non-LRCs at the base of the rete ridge, and yellow arrows mark LRCs near the top of the inter-ridge (G-I). The dashed line delineates the undulating epithelial-mesenchymal boundary. Green, H2B-GFP; Blue, Hoechst (nucleus). Scale bars: 100 μm. **(J-L)** Quantification of GFP+ cell distribution in the ventral tongue, soft palate, and hard palate based on the distance from the base of the rete ridges. A total of 100–200 GFP+ cells were counted from sagittal sections of the oral mucosa. Data are presented as mean ± S.D. (N=3). **(M-O)** Whole-mount staining of the ventral tongue, soft palate, and hard palate following 3-week chase (M, N) and 2-week chase (O). Representative H2B-GFP dilution patterns are shown. Green, H2B-GFP; Red, K14 (basal layer); Blue, Hoechst (nucleus). Scale bar: 100 μm.

To further examine the three-dimensional localization pattern of LRCs, we performed whole-mount staining of the oral epithelial layer. At the 0-day chase, we observed variations in the shape and size of the undulating structures depending on the mucosal region: a band-like pattern was observed in the ventral tongue and hard palate, while the soft palate displayed both band-like and island-like undulations (Fig. S2A-C; identified with white two-headed arrows and asterisks). Despite these structural differences, LRCs consistently localized at the inter-ridges and non-LRCs at the bottom of the rete ridges across all examined tissues (Fig. 1M-O, Fig. S2). These results demonstrate a consistent spatial segregation of slow-and fast-cycling cells along the undulating structure of the mouse oral epithelium *in vivo*.

### Dlx1+ slow-cycling and Slc1a3+ fast-cycling populations both function as stem cells in the oral mucosa

To investigate the stem cell potential and lineage relationships of slow-and fast-cycling epithelial populations, we performed lineage tracing using markers previously identified in mouse skin (Fig. 2A)^5^. We hypothesized that if only slow-cycling cells (LRCs) function as stem cells, consistent with a hierarchical stem/progenitor model^20^, Dlx1-CreER+ slow-cycling clones would contribute to the entire tissue, while Slc1a3+ fast-cycling clones would act as short-lived progenitors and eventually be lost (Fig. 2B). Conversely, if both populations possess self-renewal capacity and the ability to generate differentiated cells, their clones would persist over time in distinct regions (Fig. 2C).

**Figure 2.**
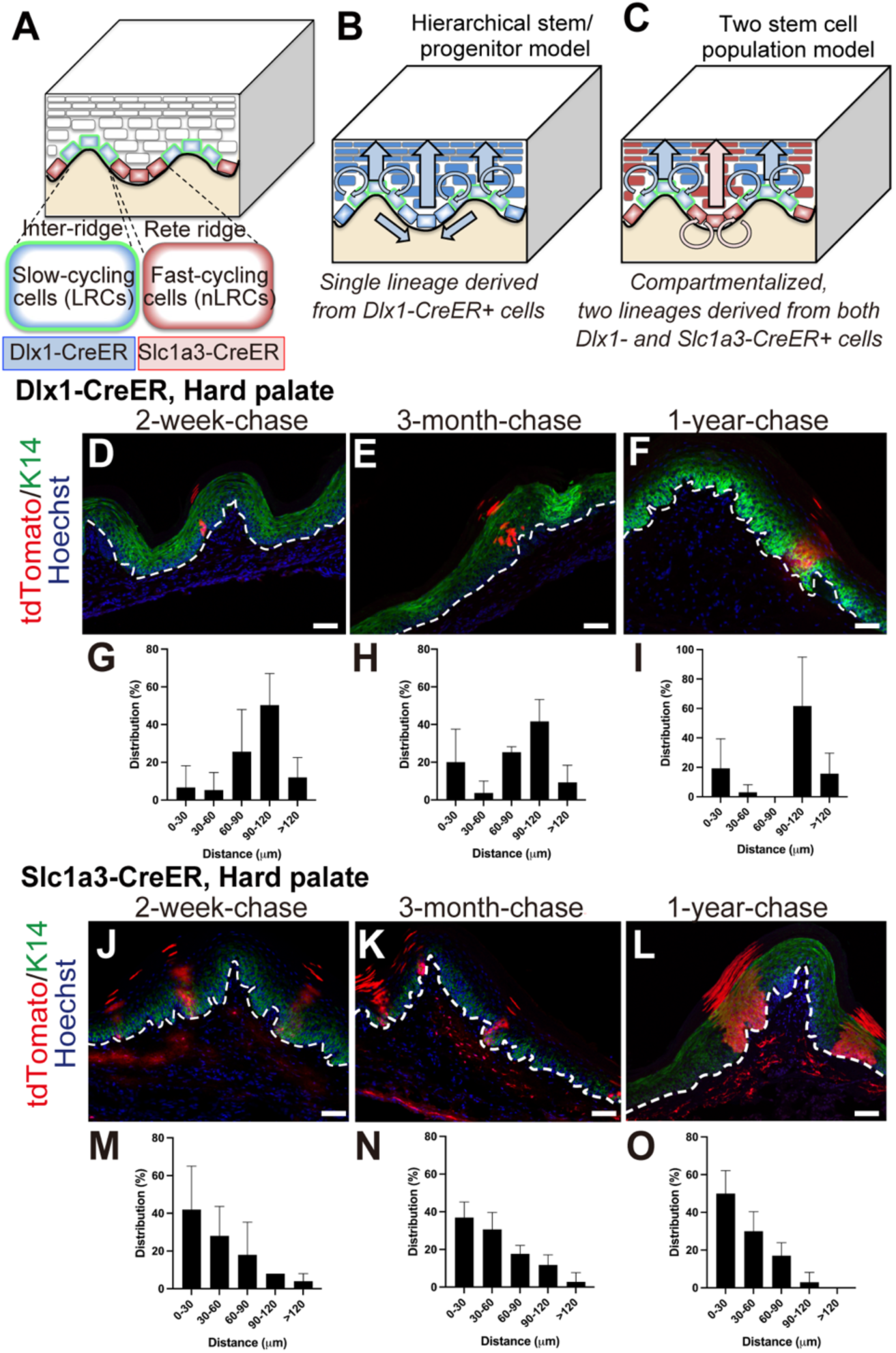
Lineage tracing with Dlx1-CreER and Slc1a3-CreER markers in the hard palate. (A-C) Models of stem cell behavior relative to the undulating structure. A single lineage derived from Dlx1-CreER+ stem cells, supporting the hierarchical stem/progenitor model (B). Two distinct lineages derived from Dlx1-CreER+ and Slc1a3-CreER+ stem cells, supporting a compartmentalized model with two stem cell populations (C). **(D-F)** Immunostaining of hard palate sections in Dlx1-CreER mice at 2-week-(D), 3-month (E), and 1-year (F) chases. Red, tdTomato; Green, K14 (basal layer); Blue, Hoechst (nucleus). The dashed line delineates the undulating epithelial-mesenchymal boundary. Scale bars: 100 μm. **(G-I)** Quantification of tdTomato+ clone distribution in Dlx1-CreER mice presented as the percentage of total clones at each chase period. A total of 50–60 tdTomato+ clones from the hard palate sections were analyzed based on their distance from the rete ridge. Data are presented as mean ± S.D. (N=3). **(J-L)** Immunostaining of hard palate sections in Slc1a3-CreER mice at 2-week (J), 3-month (K), and 1-year (L) chases. Red, tdTomato; Green, K14 (basal layer); Blue, Hoechst (nucleus). The dashed line outlines the undulating epithelial-mesenchymal boundary. Scale bars: 100 μm. **(M-O)** Quantification of tdTomato+ clone distribution in Slc1a3-CreER mice presented as the percentage of total clones at each chase period. A total of 50–60 tdTomato+ clones from the hard palate sections were analyzed based on their distance from the rete ridge. Data are presented as mean ± S.D. (N=3).

Lineage tracing with Dlx1-CreER in the hard palate showed that Tomato+ cells were preferentially located in the basal layer of the inter-ridge region (Fig. 2D-F), consistent with the distribution of H2B-GFP LRCs (Fig. 1I). Over a year, these clones expanded upwards and differentiated while remaining confined to their original inter-ridge region (Fig. 2E, F). Quantitative analysis of labeled clone distribution showed that after 2 weeks, most Dlx1-CreER+ clones were located more than 60 μm from the rete ridge base, peaking between 90 and 120 μm (Fig. 2G). After a year, the clones persisted in the 90 to 120 μm region, supporting their long-term self-renewal capacity in the inter-ridge region (Fig. 2H, I). Throughout the chase periods, there was no apparent migration of Dlx1-CreER+ clones from the inter-ridge toward the rete ridge. Instead, they replenished their lineage within the inter-ridge compartment, supporting the compartmentalized stem cell model (Fig. 2C).

In contrast, Slc1a3-CreER+ clones were primarily located in the rete ridge region, where non-LRCs reside (Fig. 2J-L). After a 2-week chase, Tomato+ cells were observed in both the suprabasal and basal layers (Fig. 2J), indicating faster division and differentiation compared to the Dlx1-CreER+ population (Fig. 2D). Quantification revealed that 40–60% of Slc1a3-CreER+ clones were within 0–30 μm of the rete ridge base after 2 weeks (Fig. 2M). Over 3 months to a year, these clones maintained their position and expanded vertically, with 50–60% remaining within the 0–30 μm region, displaying a spatially restricted pattern distinct from Dlx1-CreER+ clones (Fig. 2K, L, N, O). Similar patterns were observed in the ventral tongue and soft palate (Fig. S3, S4). Together, these findings support a compartmentalized model with two distinct stem cell populations, marked by Dlx1-CreER and Slc1a3-CreER, respectively replenishing each compartment of the inter-ridge and rete ridge regions.

### Molecularly distinct characteristics of slow-and fast-cycling populations in the skin and oral mucosa

To explore the molecular characteristics of slow-and fast-cycling populations across different epithelial tissues, we performed RNA-seq of basal LRCs and non-LRCs sorted from mouse skin and oral mucosa. K5-tTA/pTRE-H2B-GFP mice were treated with doxycycline for two weeks, and basal cell populations were isolated by sorting α6-integrin^high^/CD34^-^ clusters from the mouse back skin, tongue, and palate (Fig. S5A, D, G, J). Based on previous studies in the back skin epidermis^5^, cells with a GFP dilution rate of 1 to 3 were defined as LRCs, while cells with more than six divisions were classified as non-LRCs (Fig. S5B, C). Given the higher division rate of the oral epithelium^19,21^, cells were collected from the oral mucosa after a 10-day chase and from the skin after a 2-week chase to obtain comparable proportions of LRCs and non-LRCs across tissues (Fig. S5B, C, E, F, H, I, K, L).

Clustering and heatmap analysis revealed that oral mucosa and skin samples clustered separately, reflecting distinct transcriptional identities consistent with unique tissue features. Within each tissue, LRCs and non-LRCs exhibited different molecular profiles (Fig. 3A, Table. S1). Despite these differences, certain shared expression patterns were observed across tissues (Fig. 3B). Gene ontology analysis indicated that genes involved in protein modification, axon guidance, cell cycle, and inflammation were enriched in the LRC populations. Conversely, non-LRC populations were enriched for extracellular matrix and oxidative stress gene signatures. Notably, EGFR and TGF-β signaling pathways were enriched in non-LRCs, potentially contributing to the proliferative activity of fast-cycling stem cells (Fig. S6).

**Figure 3.**
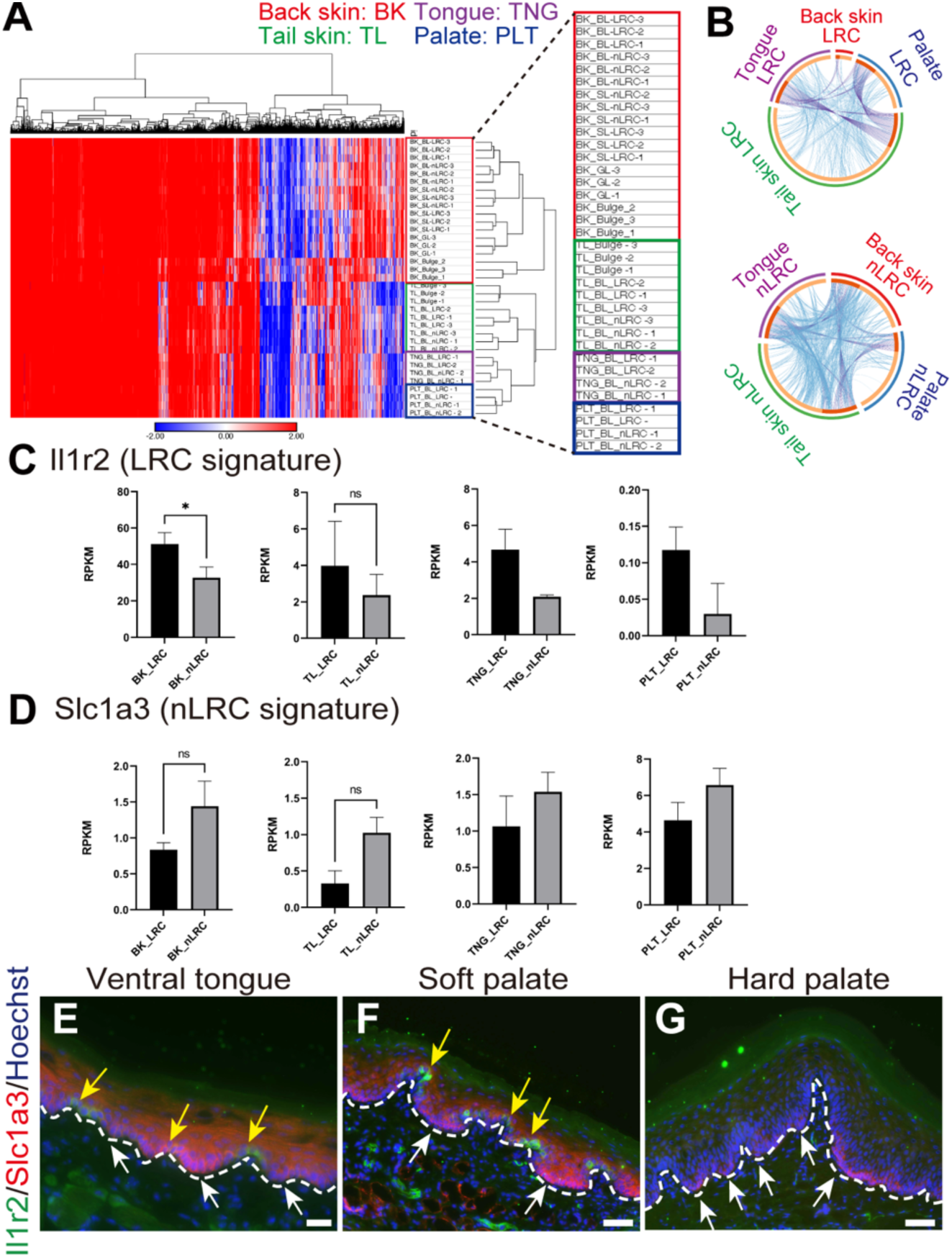
Molecular profiling of LRCs and non-LRCs in mouse skin and oral mucosa. **(A)** Hierarchical clustering of transcriptomic profiles of LRCs and non-LRCs from the mouse back skin (BK, red), tail skin (TL, green), ventral tongue (TNG, purple), and hard palate (PLT, blue). **(B)** Clustering analysis of gene expression data displayed using a circos plot. **(C, D)** Gene expression levels of *Il1r2* (an LRC signature, C) and *Slc1a3* (a non-LRC signature, D). Data are presented as mean ± S.D. Statistical significance was determined using a two-tailed t-test. BK and TL; (N=3). \**P* < 0.05. ns: not significant. TNG and PLT; (N=2; Statistical analysis not performed due to the limited sample size). **(E-G)** Immunostaining of Il1r2 and Slc1a3 in the ventral tongue (E), soft palate (F), and hard palate (G). Green, Il1r2 (LRC, yellow arrows); Red, Slc1a3 (non-LRC, white arrows); Blue, Hoechst (nucleus). The dashed line delineates the undulating epithelial-mesenchymal boundary. Scale bars: 100 μm.

To identify biomarkers for slow-and fast-cycling stem cell populations, we selected gene candidates from RNA-seq data and previous microarray datasets^5^. Among these, *Il1r2* was consistently upregulated in LRCs of both mouse skin and oral mucosa, while *Slc1a3* was enriched in non-LRCs (Fig. 3C, D). Immunostaining revealed that the Il1r2 protein localized predominantly in the inter-ridge regions of the ventral tongue, and soft palate (Fig. 3E, F; identified with yellow arrows). In contrast, the Slc1a3 protein was enriched in the rete ridge regions of the tongue, soft palate, and hard palate (Fig. 3E-G; identified with white arrows). These expression patterns correspond to the undulating structures, suggesting that Il1r2 and Slc1a3 can serve as markers for distinguishing slow-and fast-cycling stem cell populations in the oral mucosa.

### A three-dimensional culture model using a micropatterned scaffold induces proliferative heterogeneity in epidermal stem cells

To investigate the role of undulating structures in inducing stem cell heterogeneity in the epithelium, we developed an *in vitro* three-dimensional skin model using a micropatterned scaffold designed to mimic these structures (Fig. 4A, Fig. S7A)^22^. Human skin keratinocytes were seeded on the micropatterned scaffold, and differentiation and stratification were induced by the air-liquid interface to create a three-dimensional structure (Fig. S7B). This feeder-free culture system allows the direct analysis of keratinocyte phenotype and behavior in response to the undulating scaffold surface without the influence of fibroblast-derived factors. Epidermal differentiation and stratification were successfully induced on both non-patterned and micropatterned scaffolds (Fig. S7C-Q). Notably, the micropatterned scaffold significantly increased epidermal thickness, suggesting that the undulating structure may enhance the tissue-regenerative ability of epidermal stem cells (Fig. 4B, C).

**Figure 4.**
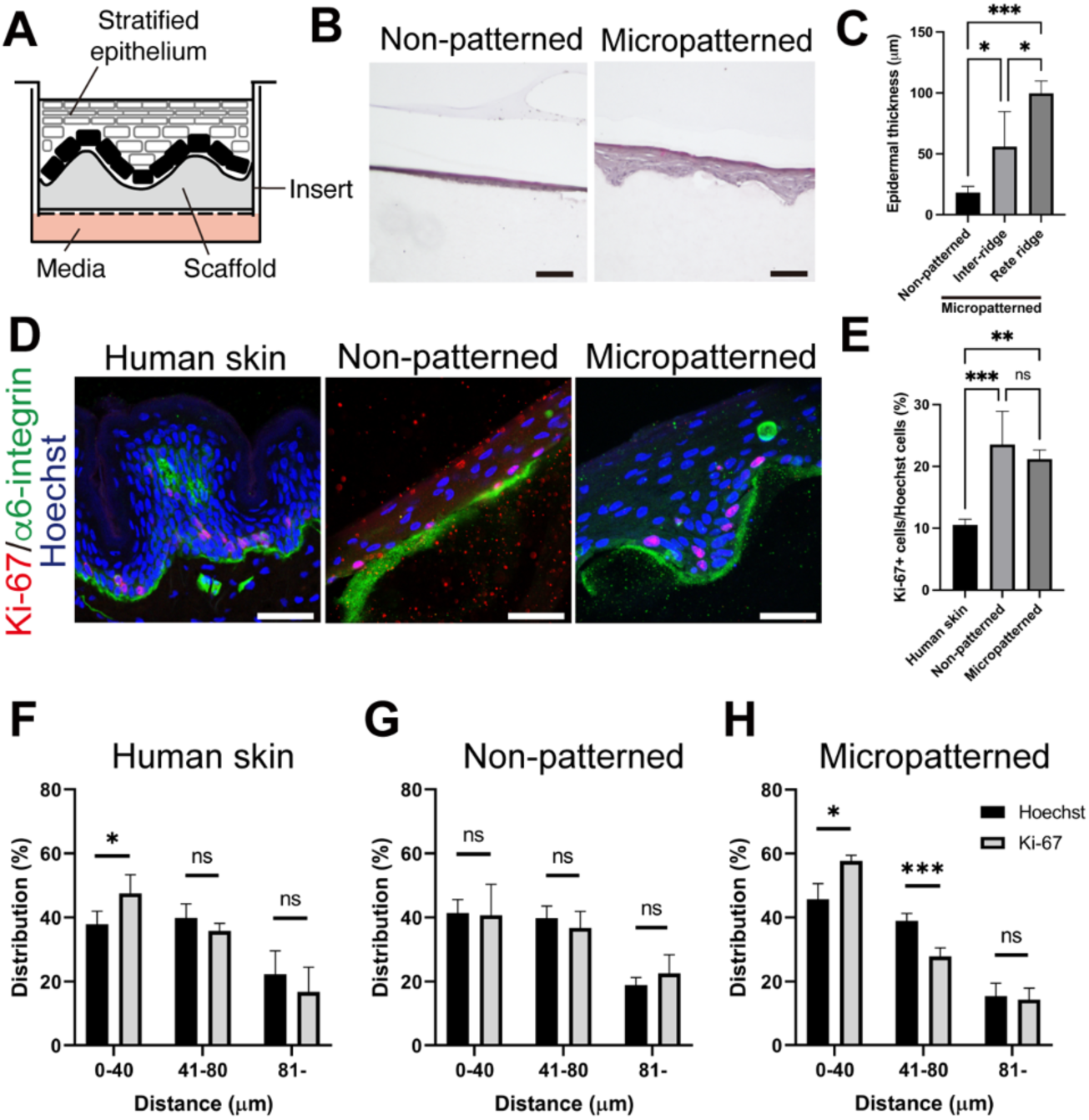
Three-dimensional skin model using a micropatterned scaffold induces proliferative heterogeneity of epidermal stem cells. **(A)** Schematic representation of the three-dimensional culture system. **(B)** Hematoxylin and eosin (H&E) staining of the skin model comparing non-patterned and micropatterned scaffolds. Scale bar: 100 μm. **(C)** Quantification of epidermal thickness. In the micropatterned scaffolds, thickness was measured from the inter-ridge or the rete ridge to the epidermal surface. Data are presented as mean ± S.D. Statistical analysis was performed using a one-way ANOVA (N=4). \*\*\**P* < 0.001; \**P* < 0.05. **(D)** Immunostaining of human skin and skin models cultured on non-patterned and micropatterned scaffolds. Red, Ki- 67; Green, α6-integrin (basement membrane); Blue, Hoechst (nucleus). Scale bars: 50 μm. **(E)** Quantification of the number of Ki-67+ proliferating cells across the conditions. Data are presented as mean ± S.D. Statistical analysis was performed using a one-way ANOVA (N=4). \*\*\**P* < 0.001; \*\**P* < 0.01. ns: not significant. **(F-H)** Distribution of Hoechst+ nuclei (black bars) and Ki-67+ cells (gray bars) in human skin (F) and skin models with non-patterned (G) and micropatterned scaffolds (H). Cell distribution was measured as the distance from the rete ridge. In the non-patterned scaffold model, the distance between the Ki-67+ cells and the edge of the structure was measured. Data are presented as mean ± S.D. Statistical analysis was performed using a two-tailed t-test (N=4). \*\*\**P* < 0.001; \**P* < 0.05. ns: not significant.

Finally, we compared the distribution of proliferating epidermal stem cells on non-patterned and micropatterned scaffolds by assessing Ki-67 expression. No significant difference was observed in the number of Ki-67+ proliferating cells between the two scaffold types (Fig. 4D, E). Both scaffolds exhibited a higher proportion of Ki-67+ cells compared to human skin, consistent with previous reports^23^. In human skin, Ki-67+ cells were predominantly localized within 0–40 μm from the base of the rete ridges, compared to all basal cells stained with Hoechst (Fig. 4F). On non-patterned scaffolds, Ki-67+ cells displayed an unbiased distribution throughout the basal layer (Fig. 4G). However, in the micropatterned scaffolds, 55–60% of Ki-67+ cells were preferentially located within 0–40 μm from the base of the rete ridges, closely mirroring the distribution observed in human skin (Fig. 4H). These findings suggest that the micropatterned scaffold plays a functional role in inducing heterogeneous division patterns of epidermal stem cells, providing potential mechanistic insight into analogous undulated structures.

## Discussion

In this study, we demonstrated that stem cells in the mouse oral mucosa are heterogeneous, comprising two distinct populations with differing cell division rates, gene expression profiles, and spatial localization patterns associated with an undulating tissue structure. The undulating architecture partially induces proliferative heterogeneity in a three-dimensional culture system designed to recapitulate these surface structures. Our findings suggest that an undulating tissue architecture serves as a potential regulator of proliferative heterogeneity, highlighting the importance of the mechanical environment as a conserved mechanism influencing stem cell heterogeneity across epithelial tissues. Additionally, our results indicate that the mouse oral mucosa, with its characteristic undulating structure, can serve as a valuable *in vivo* model for exploring the relationship between the surface topography and stem cell systems in both human skin and oral mucosa.

Using lineage tracing and H2B-GFP pulse-chase experiments, we demonstrated that both slow-and fast-cycling populations exhibit properties of oral epithelial stem cells, with their localization corresponding to the undulating structure. The undulating features described in this study, specifically rete ridges and inter-ridges, are approximately 50–100 μm in depth in the mouse oral mucosa, while the larger rugae structures of the hard palate reach depths of 300–500 μm. Our data indicate that the Dlx1-CreER and Slc1a3-CreER labeled populations are primarily localized in the junctional zone of the hard palate and likely represent a subset of the Lrig1-CreER+ population previously identified by Byrd et al^12^.

In this study, we identified Il1r2 (Interleukin 1 receptor, type II) as a marker for the slow-cycling population in the oral mucosa (Fig. 3C). Il1r2 functions as a decoy receptor for IL-1, inhibiting IL-1α and IL-1β-mediated signal transduction^24^. Although Il1r2 is primarily known for its anti-inflammatory effects, its role in epithelial stem cell regulation remains unclear. In hair follicle stem cells, prolonged IL-1 signaling induced by a high-fat diet has been shown to deplete the stem cell pool, leading to hair loss^25^. It is possible that the expression of Il1r2 in slow-cycling epithelial stem cells serves a protective role, shielding these cells from inflammatory signals and preserving the stem cell pool.

The precise mechanisms by which the undulating structure regulates tissue stem cell heterogeneity remain to be fully elucidated. However, the coordinated influence of biochemical and biomechanical factors may contribute to the formation of distinct microenvironments^26–28^. For instance, in the intestinal epithelium, epithelial bending during development concentrates Shh signaling at the villus tips, restricting intestinal stem cells to the base of the crypts^26^. Similarly, in an *in vitro* model where human keratinocytes were cultured on a PDMS scaffold mimicking undulating tissue architecture, integrin β1 and nuclear YAP were localized at the peaks of the undulations^15,29^. These studies suggest that the mechanical environment surrounding the cells may differ based on cellular positioning within undulating structures, potentially contributing to the induction of heterogeneity among epithelial stem cells.

Three-dimensional skin models have been developed for clinical and research applications^30^. Co-culture systems combining keratinocytes and fibroblasts are commonly employed to reconstruct the skin tissue^31,32^, clarifying the niche environment formed through cell-cell interactions and secreted factors. However, such systems present challenges in isolating the effects of mechanical cues generated by the undulating structure itself. Our model, featuring a micropatterned scaffold that mimics the undulating structure of the skin and oral mucosa, provides a unique platform for studying the significance of these mechanical influences while minimizing the effects of dermal cells. This culture system allows for precise manipulation of scaffold properties, such as stiffness, the geometry of undulation, and the composition of the extracellular matrix^33,34^. Consequently, this innovative three-dimensional culture model offers a tool for modulating the behavior of epidermal stem cells through scaffold modification, with potential applications in both fundamental research and clinical fields, including tissue transplantation and drug discovery.

## Acknowledgments

We would like to thank the Center for Animal Resources and Development at Kumamoto University, the Animal Resource Center at the University of Tsukuba, and the Laboratory of Embryonic and Genetic Engineering at Kyushu University for their excellent mouse care. We also thank the International Core-facility of Advanced Life Science at Kumamoto University and the Research Promotion Unit at Kyushu University for their invaluable support. We thank T. Keida (Kumamoto University) and K. Kawazoe (University of Tsukuba) for their technical assistance with immunostaining, and Dr. M. Muratani (University of Tsukuba) for his advice on RNA-seq analysis. We also thank T. Kawakami (Taki Chemical Co., Ltd.), T. Komatsu, and R. Nakamura (Komatsuseiki Kosakusho Co., Ltd) for their advice and technical support with the micropatterned scaffolds. This work was supported by the Program for Technological Innovation of Regenerative Medicine, AMED (23bm0704067) (to A.S.), AMED-PRIME, AMED (21gm6110016) (to A.S.), Interstellar Initiative Beyond, AMED (22jm0610063) (to A.S.), Grant-in-Aid for Scientific Research (B) (20H03266, 24K02035) (to A.S.), Grant-in-Aid for Early-Career Scientists (18K14709) (to A.S.), and (23K14198) to (M.I.), and Uehara Memorial Foundation (to A.S.), The Takeda Foundation (to A.S.), the Sasakawa Scientific Research Grant from The Japan Science Society (to M.I.), and the IKEDARIKA Grant from Ikedarika Scientific Co., Ltd., and Leave a Nest Co., Ltd. (to M.I.). This work was supported in part by the MEXT Cooperative Research Project Program, Medical Research Center Initiative for High Depth Omics, and CURE: JPMXP1323015486 for MIB, Kyushu University. Additionally, we acknowledge support from JST SPRING, Grant Numbers JPMJSP2124 (to Y.X.N) and JPMJSP2136 (to I.N.).

## Author Contributions

A.S. conceptualized the project, designed the experiments, and contributed expertise in stem cell and skin biology. A.S. and H.Y. managed and supervised the project. M.I., Y.X.N., I.N., A.S., H.K., and K.I. performed experiments and analyzed the results. K.I., H.K., R. Maeda, R. Mizuno, and J.M. provided insights into oral mucosal analysis and developed the micropatterned scaffold. H.Y. contributed knowledge regarding the extracellular matrix and mechanobiology. The manuscript was drafted by M.I. and Y.X.N. and edited by A.S. Funding was acquired by A.S. and M.I.

## Declaration of Interests

K.I. is the CEO of CollaWind Co. Ltd. J.M. is the CTO of CollaWind Co. Ltd. M.I. and A.S. received donations of micropatterned and non-patterned scaffolds from CollaWind Co. Ltd. (Niigata, Japan).

## Declaration of generative AI and AI-assisted technologies in the writing process

During the preparation of this manuscript, the authors used DeepL, Grammarly, and ChatGPT to assist with language clarity and readability. Following the use of these tools, the authors thoroughly reviewed and edited the content to ensure accuracy and take full responsibility for the final version of the published article.

## Materials and Methods

### Animals and Ethics

All animal procedures complied with the guidelines for animal experimentation approved by the Institutional Animal Experiment Committee at Kumamoto University, the University of Tsukuba, and Kyushu University. K5-tTA^35^ (a gift from Dr. Adam Glick)/pTRE-H2B-GFP^17^ (The Jackson Laboratory, no. 005104) transgenic were used for H2B-GFP dilution experiments and the isolation of LRCs and non-LRCs. For lineage tracing, Dlx1-CreER mice^36^ (The Jackson Laboratory, no. 014551) and Slc1a3-CreER mice (The Jackson Laboratory, no. 012586) were crossed with Rosa-tdTomato reporter mice (The Jackson Laboratory, no. 007905). Both male and female mice were used in all experiments. Experimental mice were housed in the Center for Animal Resources and Development at Kumamoto University, the Laboratory Animal Resource Center at the University of Tsukuba, and the Laboratory of Embryonic and Genetic Engineering at Kyushu University.

### Doxycycline and Tamoxifen Treatment

For H2B-GFP pulse-chase experiments, mice were fed with doxycycline chow (1 g/kg; Oriental Kobo Inc.) for the indicated chase periods, beginning at 2–3 months of age. GFP expression was detected using anti-GFP antibody staining after 0-day, 2-week, and 3-week chase periods. For lineage tracing of slow-cycling (Dlx1-CreER) and fast-cycling (Slc1a3-CreER) cells, mice were injected intraperitoneally with tamoxifen (100 μg/g body weight; Sigma-Aldrich) for five consecutive days at 2–3 months of age. Mice were sacrificed for analysis at 2-week, 3-month, and 1-year chase periods following tamoxifen administration.

### Whole-Mount Immunostaining

To separate the epithelium from the underlying connective tissues as an intact sheet, tongues, and palates were bisected and incubated in EDTA (20 mM; DOJINDO Laboratories) diluted in PBS on an orbital shaker at 37 ℃ for three hours. The epithelial sheets were fixed in 4% paraformaldehyde (PFA; FUJIFILM Wako Pure Chemical Corporation) overnight at 4 ℃ with gentle shaking. Following fixation, the epithelial sheets were washed with PBS and incubated for 3 hours at room temperature in blocking buffer containing 1% bovine serum albumin (Merck), 2.5% normal donkey serum (Sigma-Aldrich), 2.5% normal goat serum (Sigma-Aldrich), and 0.8% Triton X-100 (FUJIFILM Wako Pure Chemical Corporation) in PBS. The sheets were then incubated overnight at room temperature with primary antibodies diluted in blocking buffer. After primary antibody incubation, samples were washed four times in 0.2% Tween (Sigma-Aldrich) in PBS for 1 hour at room temperature. Secondary antibody incubation was performed overnight at 4 ℃ After washing, samples were counterstained with Hoechst (Sigma-Aldrich) for 1 hour and mounted for imaging. The primary antibodies and dilutions used were: rabbit anti-K14 (1:1000; BioLegend, 905304) and chicken anti-GFP (1:500; Abcam, ab13970). Secondary antibodies (Alexa 488, 546, or 647, Invitrogen) were used at 1:200–1:500 dilutions. Samples were mounted in antifade mounting medium and imaged using a confocal microscope (Nikon, A1 HD25 and Zeiss, LSM 900).

### H&E Staining and Section immunostaining

For histological analysis, oral mucosa and skin samples were snap-frozen in OCT compound (Tissue-Tek, Sakura Finetek). Cryosections (10 μm) were air-dried, fixed in 4% PFA for 10 min at room temperature, and washed with PBS. Sections were stained with Hematoxylin (Wako) for 20 min and Eosin Y (Wako) for 5 min. After dehydration, the sections were mounted using Entellan new mounting medium (Merck Millipore). Images were acquired using an EVOS M5000 Imaging system (ThermoFisher Scientific).

For immunofluorescence staining, 10-µM frozen sections were fixed in 4% PFA for 10 min at room temperature, followed by three 5-minute washes in 20 mM glycine (FUJIFILM Wako Pure Chemical Corporation) in PBS. Sections were then permeabilized with 0.1% Triton X-100 in PBS for 5 min. Samples were blocked for 1 hour at room temperature in PBS containing 1% bovine serum albumin (BSA; Merck), 2.5% normal goat serum (Sigma-Aldrich), 2.5% normal donkey serum (Sigma-Aldrich), 2% gelatin (Sigma-Aldrich), and 0.1% Triton X-100. Sections were incubated with primary antibodies for 1 hour at room temperature. The following primary antibodies and dilutions were used: chicken anti-GFP (1:500; Abcam, ab13970), rabbit-anti K14 (1:1000; BioLegend, 905304), rat anti-Il1r2-BV421 (1:100; BD, 562926), guinea pig anti-Slc1a3 (1:100; Frontier Institute, GLAST-GP-Af1000), mouse anti-Ki67 (1:200; BD Bioscience, 14-5698-82), rat anti-α6-integrin (1:500; BD Bioscience, 555734), chicken anti-K5 (1:500; BioLegend, 905904), mouse anti-K10 (1:500; Dako, M7002), and rabbit anti-Loricrin (1:100; Abcam, ab85679). Following primary incubation, sections were washed three times in 0.1% Triton X-100 in PBS and incubated with species-appropriate secondary antibodies conjugated to Alexa Fluor 488, 555, or 647 (1:300; ThermoFisher Scientific) for 1 hour at room temperature. Before being mounted, sections were counterstained with Hoechst (Sigma-Aldrich) for 10 min. Sections were then mounted using antifade mounting medium and imaged using a confocal microscope (Nikon, A1 HD25 and Zeiss, LSM 900).

### Fluorescence-Activated Cell Sorting (FACS)

Skin and oral tissues, including back skin, tail skin, hard palate, and ventral tongue, were dissected from K5-tTA/pTRE-H2B-GFP mice. Tissues were incubated overnight at 4 ℃ in 0.25% trypsin (MP Biomedicals) with EDTA, followed by an additional 30 min incubation at 37 ℃ the next day. Blood, fat, and connective tissues were removed by scraping. Single-cell suspensions of epithelial cells were obtained by mechanical scraping followed by filtration through a cell strainer. For cell surface marker labeling, the following antibodies and dilutions were used to incubate the cells for 30 min on ice: Streptavidin-APC (1:100; BD Biosciences, 554067), α6-integrin-BUV395 (1:200; BD Biosciences, 563271), and CD34-biotin (1:50; eBioscience, 13-0341-85). The FACS Aria flow cytometer (BD Biosciences) was used to sort the LRC and non-LRC populations. FlowJo software (BD Biosciences) was used to analyze the data.

### RNA-Sequencing Analysis

Basal cells were sorted from the back skin, tail skin, tongue, and palate of K5-tTA/pTRE-H2B-GFP mice following a 10-day chase (tongue and palate) or a 2-week chase (back and tail skin). FACS-sorted cells were directly lysed in TRIzol reagent (Ambion) and submitted to Tsukuba i-Laboratory LLP, the University of Tsukuba, for RNA extraction and sequencing. RNA-seq libraries were prepared using the SMARTer® Stranded Total RNA-Seq Kit v2 – Pico Input Mammalian (Takara) according to the manufacturer’s protocol. Sequencing was performed on an Illumina NextSeq500 platform. Raw sequencing data were analyzed using CLC Genomics Workbench 11 (Qiagen). Genes with zero expression were excluded from the dataset. Following normalization, genes with at least a 2-fold upregulation or downregulation were selected for further analysis. Hierarchical clustering and principal component analysis (PCA) were performed using the online tools Morpheus (https://software.broadinstitute.org/morpheus), Clustvis, Panther, Metascape, and VENNY 2.1 (https://bioinfogp.cnb.csic.es/tools/venny/index.html). Gene ontology and pathway analysis were conducted using g:Profiler. RNA-seq data have been deposited in the Gene expression Omnibus (GEO) under accession number GSE208730 (https://www.ncbi.nlm.nih.gov/geo/query/acc.cgi?acc=GSE208730).

### Three-Dimensional Skin Model with a Micropatterned Scaffold

Micropatterned and non-patterned collagen scaffolds derived from fish scales (CollaWind Sheet) were obtained from CollaWind Co. Ltd. (Niigata, Japan). The micropatterned scaffold dimensions were designed with a depth of approximately 100 µm and a channel width of 169 µm^22^.

Neonatal primary human keratinocytes (BIOPREDIC International) were cultured in 100 mm dishes coated with collagen type I (Corning) using CnT-Prime Epithelial Culture Medium (CELLnTEC) until reaching 80–90% confluency. Non-patterned or micropatterned scaffolds were pre-coated with 500 µL of collagen type IV (1mg/mL; Sigma-Aldrich) overnight at 4 ℃ The following day, a keratinocyte cell suspension (5 x 10^5^ cells/mL in CnT-PR medium) was seeded onto the scaffolds at a density of 6.7 x 10^5^ cells per scaffold. After seeding, cells were cultured in CnT-PR medium and incubated at 37 ℃ in 5% CO2. Once confluency was reached, the culture medium was switched from CnT-PR to CnT-Prime Epithelial 3D Airlift Medium (CELLnTEC) and incubated for 15–16 hours. Subsequently, the medium was aspirated, and the scaffolds were transferred to a 6-well deep plate (Corning, 355467) containing a culture insert (Falcon, 353091). Fresh CnT-PR-3D medium (∼10 mL) was added to reach the membrane surface, initiating the air-liquid interface (ALI) culture. The ALI culture was maintained for 8 days, with CnT-PR-3D medium replaced every 2 days. Following the ALI culture, samples were collected and prefixed in 4% PFA at room temperature for 2 hours. Prefixed samples were then embedded in 4% agarose (Ina Food Industry Co., Ltd.) and subsequently soaked in a 30% sucrose/PBS solution at room temperature for 16 to 17 hours. Samples were then embedded in OCT compound.

### Human Skin Samples

Frozen full-thickness normal human abdominal skin samples at ages of 20-40s were purchased from CTI-Biotech (Lyon, France) under ethical considerations. Informed consent was obtained from anonymous donor patients for the collection of these samples, and the consent and collection procedures were compliant with European standards and applicable local ethical guidelines.

### Quantification and Statistical Analysis

Quantifications using mouse samples were conducted independently on tissue sections from more than three mice. Image analysis and measurements were conducted using ImageJ software (version 1.54f, NIH). To quantify the distribution of LRCs in the mouse oral mucosa, Ki-67+ cells in human skin and skin models, and Hoechst-stained nuclei, the distance between the bottom of the rete ridges and the target cells was measured. For the skin model with a non-patterned scaffold, which lacks an undulating structure, the distance between the bottom of the rete ridge and the next inter-ridge was determined using a micropatterned scaffold, and the average distance was used to define a single structural unit for measurements. Statistical analyses were performed using GraphPad Prism 9. Data are presented as mean ± standard deviation (S.D.).

## Supplemental Information

**Supplementary Figure 1.**
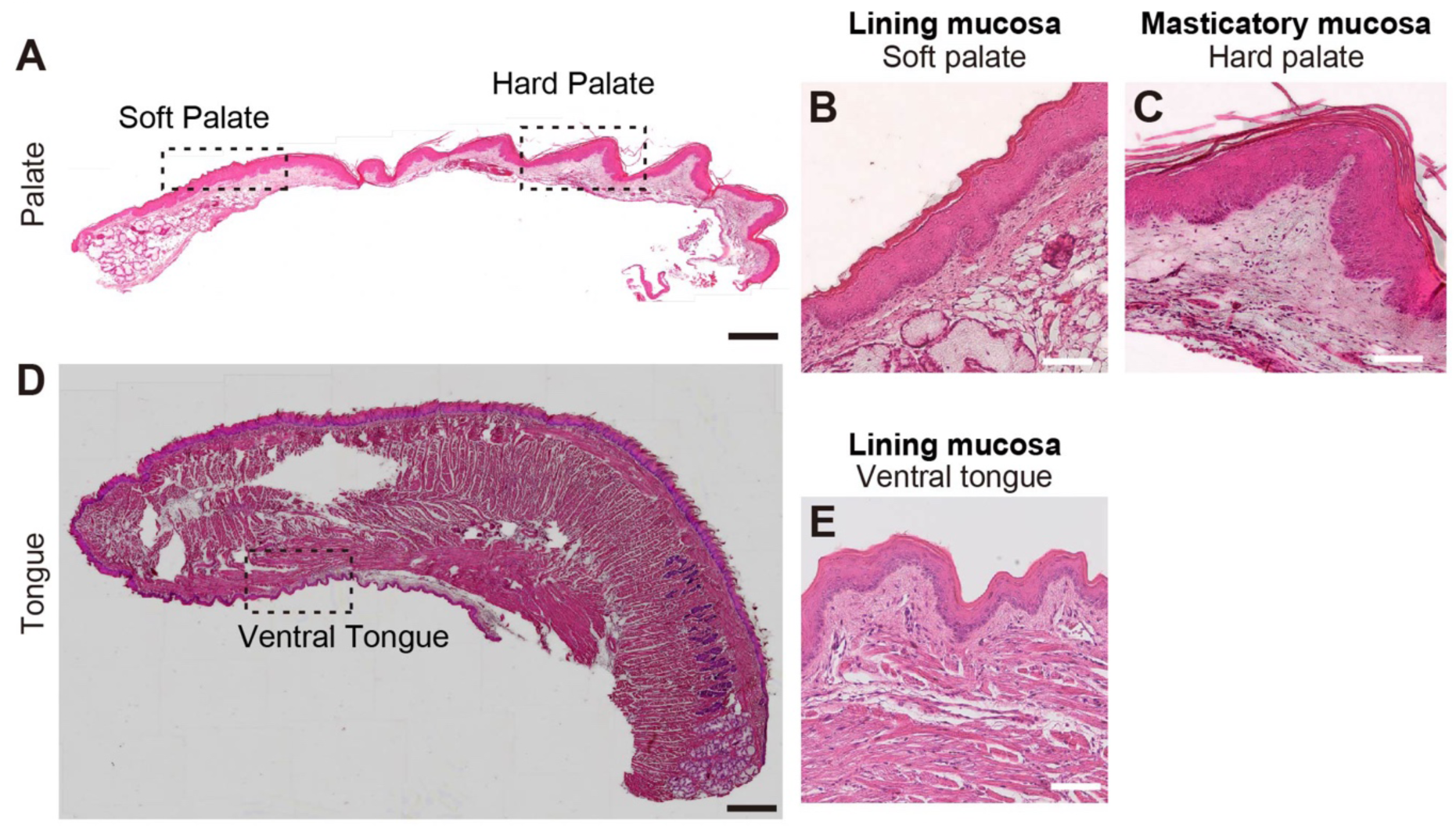
H&E staining of mouse oral mucosa. (A-E) H&E staining images of the palate (A) and tongue (D). Lining mucosa: soft palate (B), ventral tongue (E). Masticatory mucosa: hard palate (C). Scale bars: 500 μm (A, D); 100 μm (B, C, E).

**Supplementary Figure 2.**
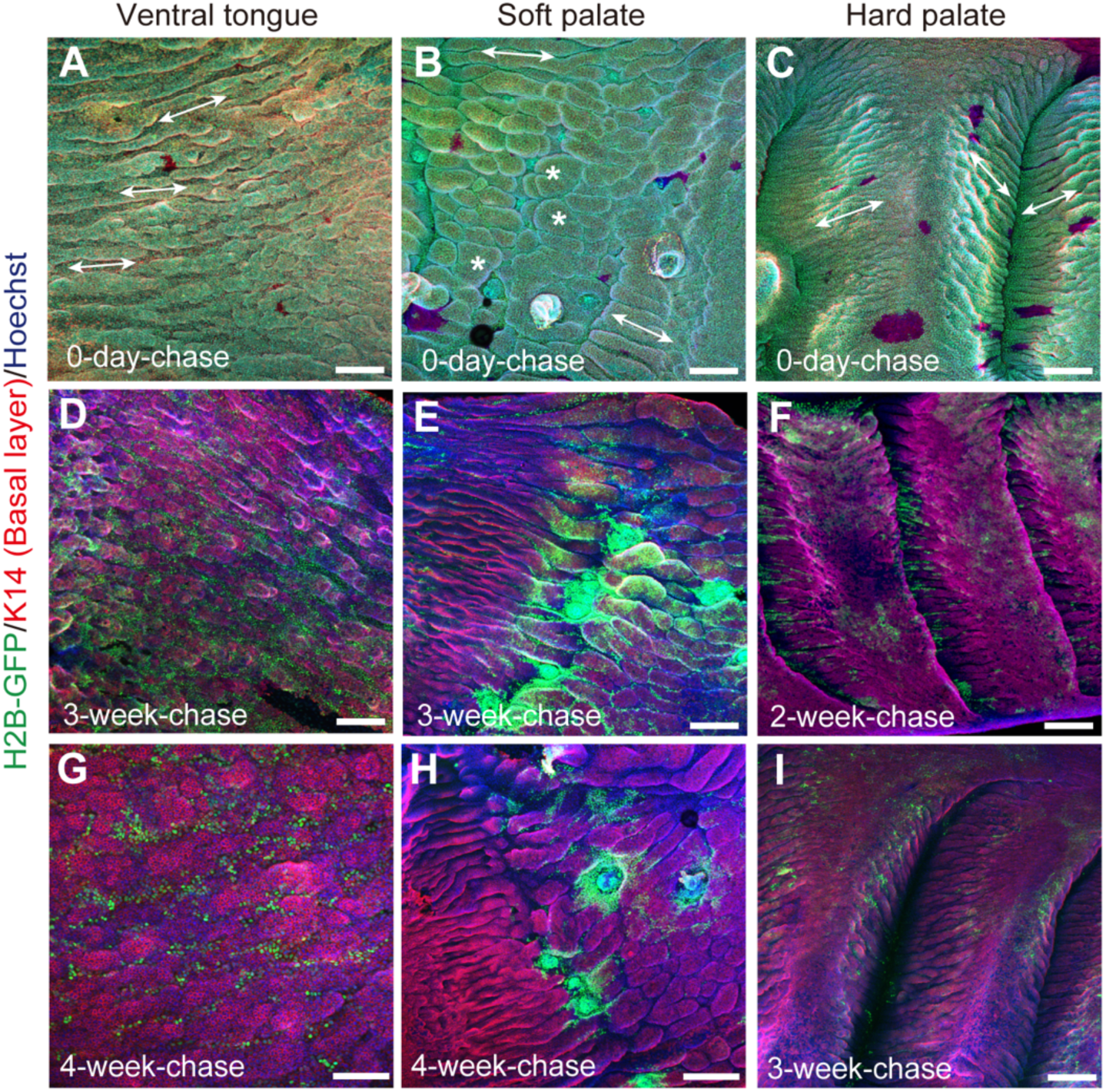
H2B-GFP label retention assay in the ventral tongue, soft palate, and hard palate. (A-I) Whole-mount immunostaining of ventral tongue, soft palate, and hard palate at 0-day, 2-or 3-week-, and 3-or 4-week chases. Green, H2B-GFP; Red, K14 (basal layer); Blue, Hoechst (nucleus). White double-headed arrows indicate band-like undulations (A-C), and white asterisks indicate island-like undulations (B). Scale bars: 200 μm (A-F, H, I); 100 μm (G).

**Supplementary Figure 3.**
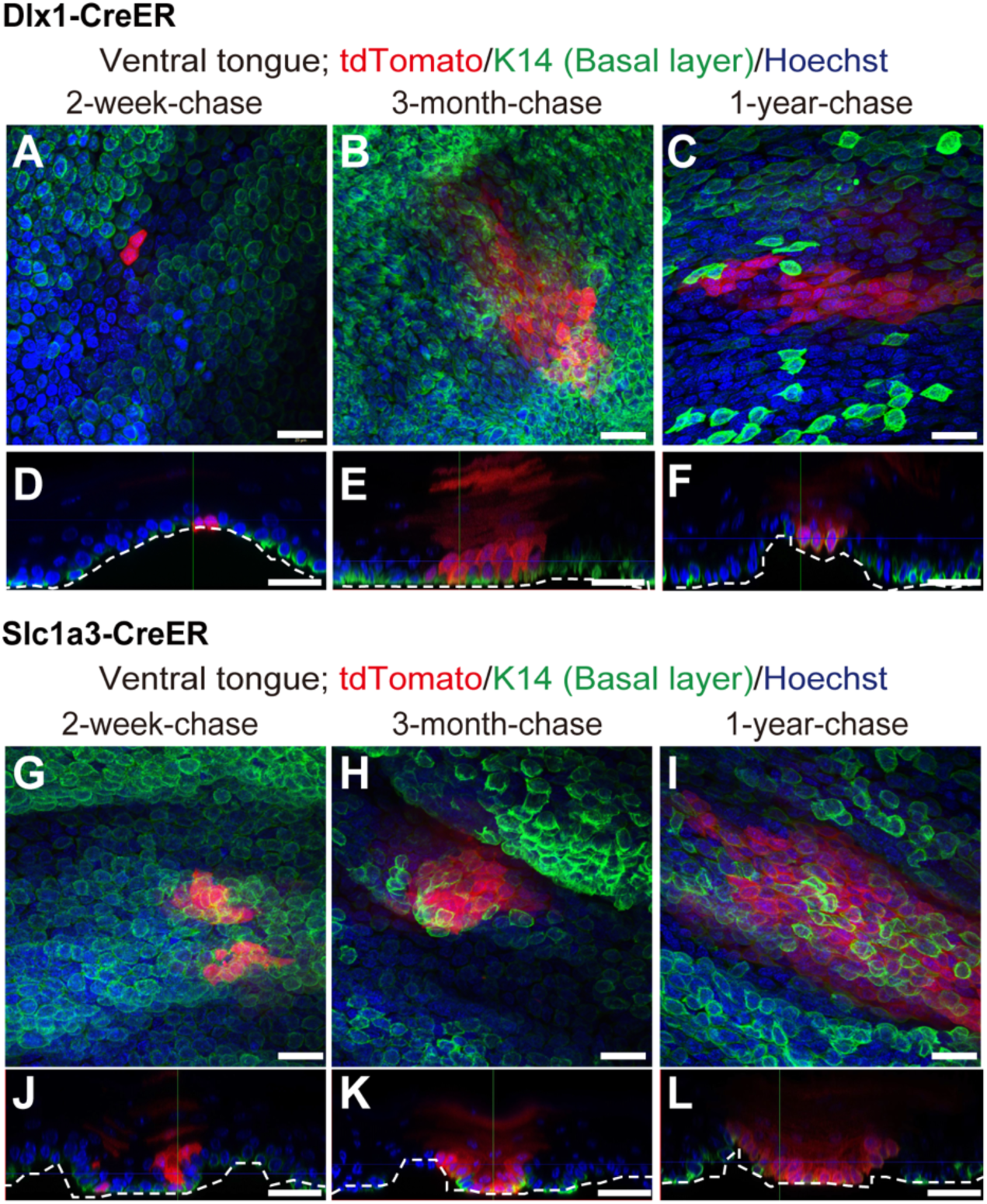
Lineage tracing with Dlx1-CreER and Slc1a3-CreER markers in the ventral tongue. (A-F) Whole-mount staining and Z-stack images of representative Dlx1-CreER+ clones at 2-week (A, D), 3-month (B, E), and 1-year (C, F) chases. **(G-L)** Whole-mount staining and Z-stack images of representative Slc1a3-CreER+ clones at 2-week (G, J), 3-month (H, K), and 1-year (I, L) chases. Red, tdTomato; Green, K14 (basal layer); Blue, Hoechst (nucleus). The dashed line outlines the undulating epithelial-mesenchymal boundary. Scale bars: 20 μm.

**Supplementary Figure 4.**
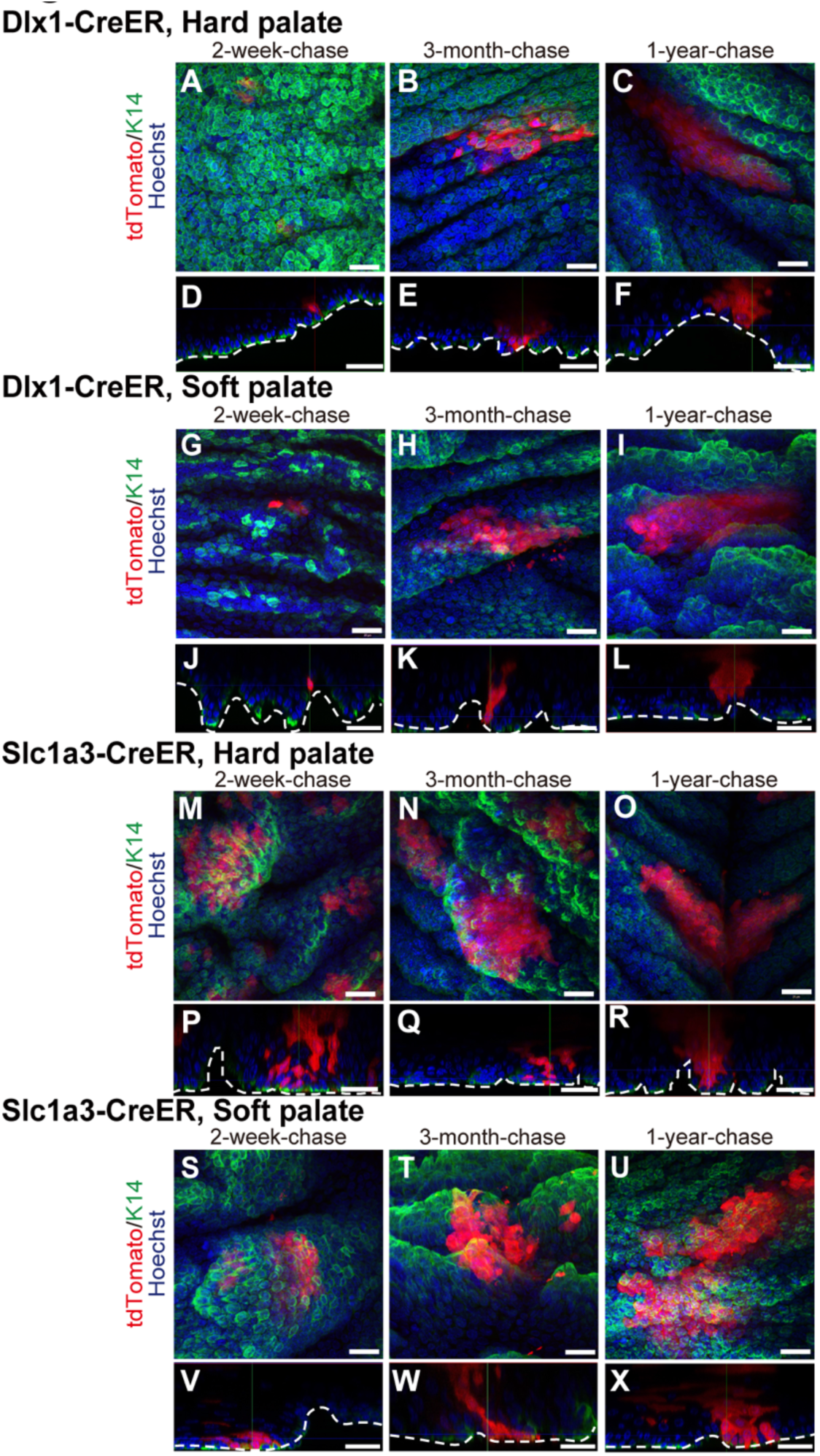
Lineage tracing with Dlx1-CreER and Slc1a3-CreER markers in the hard and soft palate. (A-F) Whole-mount staining and Z-stack images of representative Dlx1-CreER+ clones at 2-week (A, D), 3-month (B, E), and 1-year (C, F) chases in the hard palate. **(G-L)** Whole-mount staining and Z-stack images of representative Dlx1-CreER+ clones at 2-week (G, J), 3-month (H, K), and 1-year (I, L) chases in the soft palate. **(M-R)** Whole-mount staining and Z-stack images of representative Slc1a3-CreER+ clones at 2-week (M, P), 3-month (N, Q) and 1- year (O, R) chases in the hard palate. **(S-X)** Whole-mount staining and Z-stack images of representative Dlx1-CreER+ clones at 2-week (S, V), 3-month (T, W), and 1-year (U, X) chases in the soft palate. Red, tdTomato; Green, K14 (basal layer); Blue, Hoechst (nucleus). The dashed line outlines the undulating epithelial-mesenchymal boundary. Scale bars: 20 μm.

**Supplementary Figure 5.**
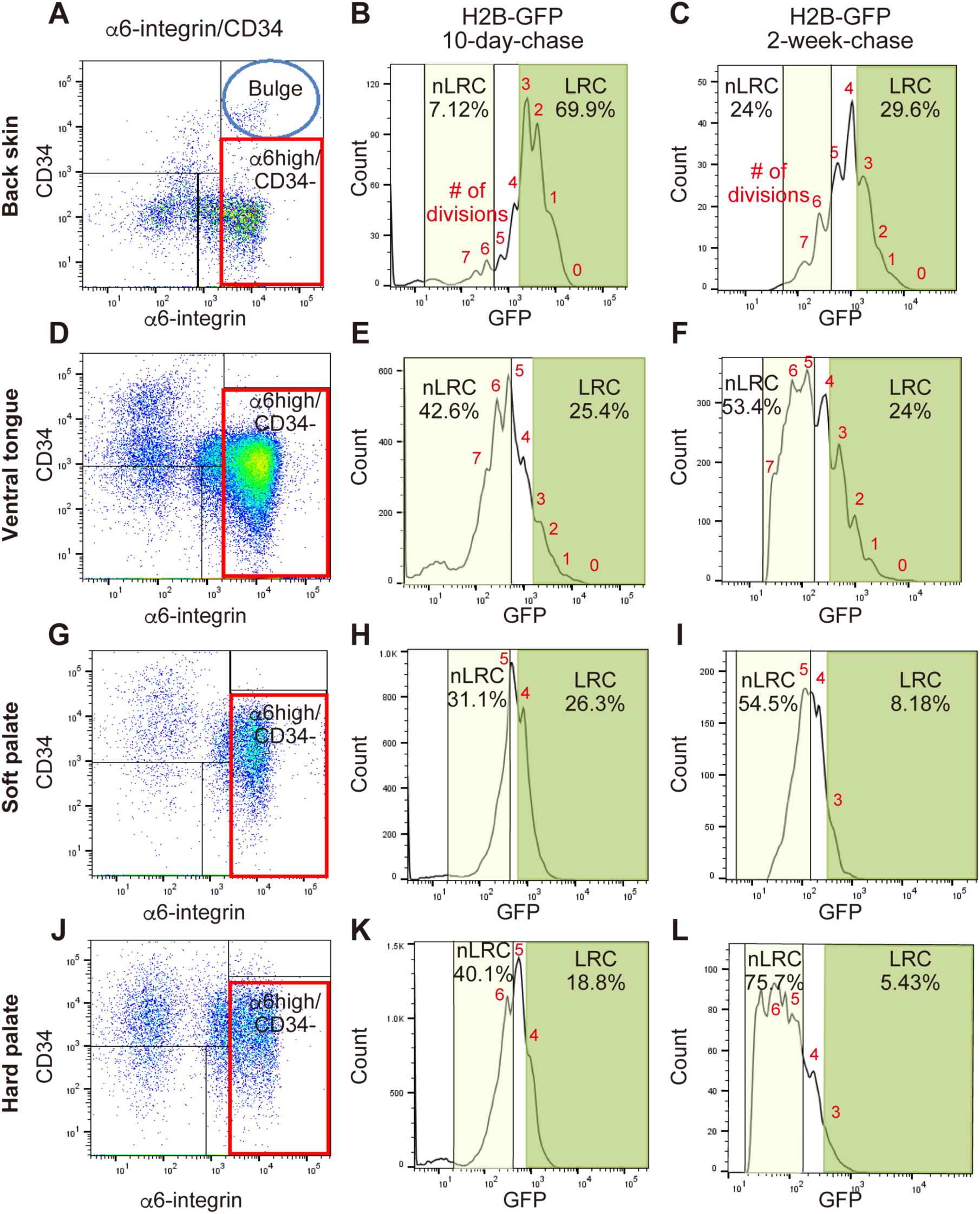
FACS analysis of cell division dynamics in the mouse skin and oral mucosa. (A-L) Representative FACS plots. Scatter plots show α6-integrin and CD34 staining. H2B-GFP fluorescence intensity in basal cell populations (α6-integrin+/CD34-) is shown as a histogram. The number of cell divisions is indicated in red.

**Supplementary Figure 6.**
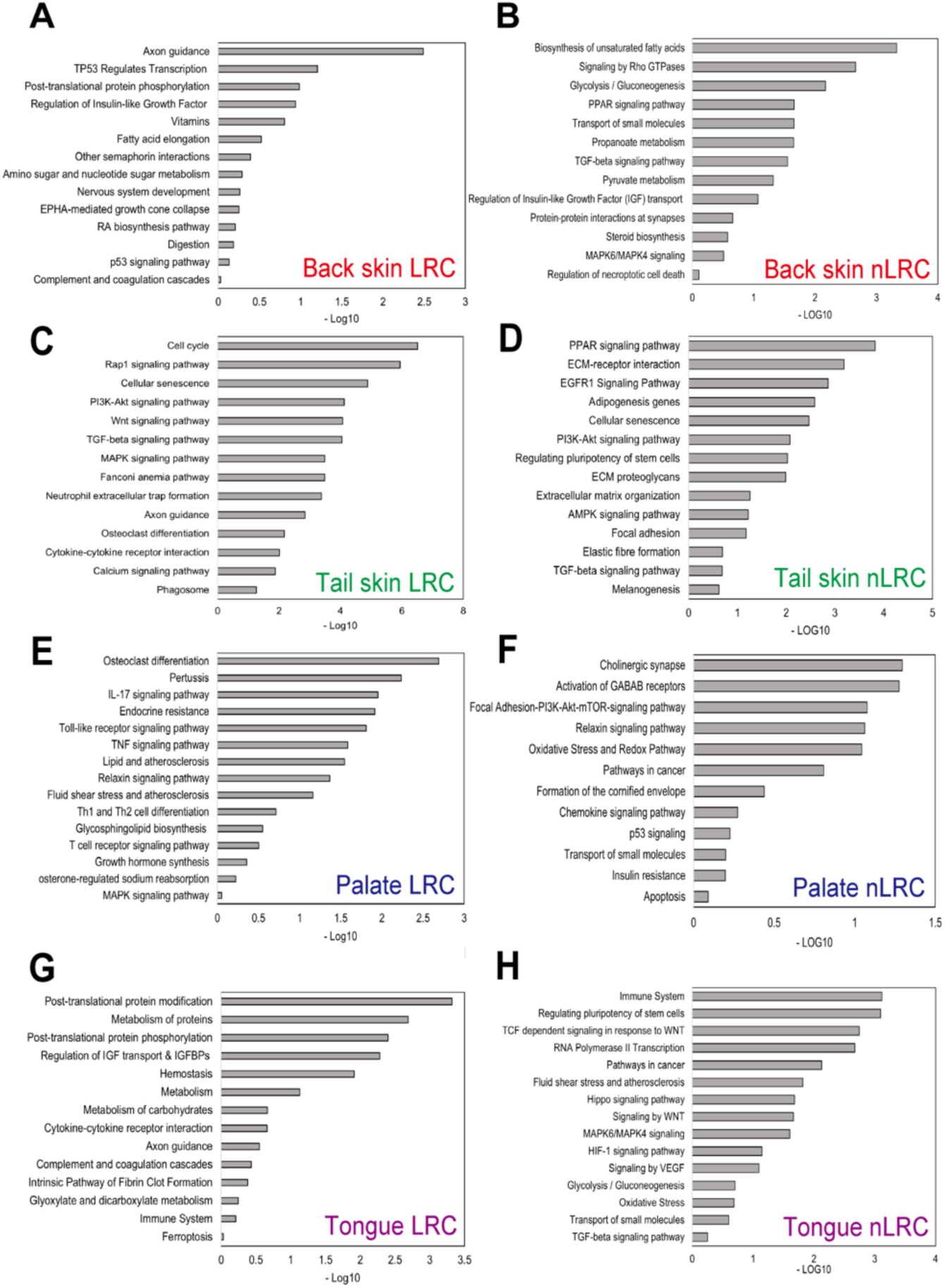
Pathway analysis of LRC and non-LRC signature genes in skin and oral mucosa. (A-H) Gene ontology analysis of LRC and non-LRC signatures in the back skin (A, B), tail skin (C, D), hard palate (E, F), and ventral tongue (G, H).

**Supplementary Figure 7.**
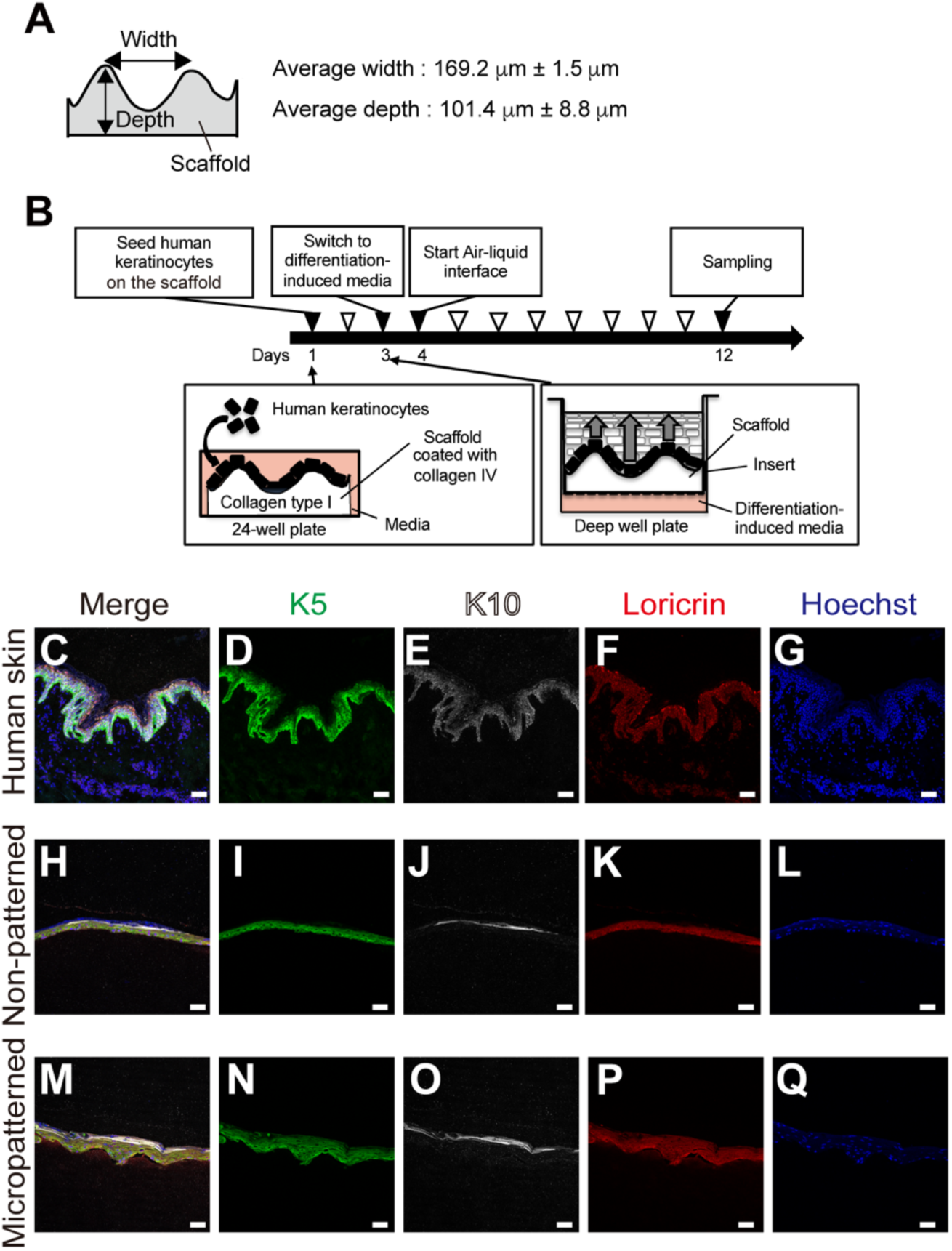
Design and analysis of three-dimensional skin models. **(A)** Schematic representation of the micropatterned scaffold used in this study. **(B)** Experimental design for the three-dimensional skin culture system. **(C-Q)** Immunofluorescence staining for epidermal basal and differentiation markers in human skin and skin models. Green, K5 (basal cell marker); White, K10 (the early-stage differentiation marker); Red, Loricrin (the late-stage differentiation marker), and Blue, Hoechst (nucleus). Scale bars: 50 μm.

**Supplementary Table 1. Signature genes of LRCs and non-LRCs in the back skin, tail skin, palate, and tongue**

## References

1. Barker, N., van Es, J.H., Kuipers, J., Kujala, P., van den Born, M., Cozijnsen, M., Haegebarth, A., Korving, J., Begthel, H., Peters, P.J., et al. (2007). Identification of stem cells in small intestine and colon by marker gene Lgr5. Nature 449, 1003–1007. 10.1038/nature06196.

2. Ishii, R., Yanagisawa, H., and Sada, A. (2020). Defining compartmentalized stem cell populations with distinct cell division dynamics in the ocular surface epithelium. Development 147, dev197590. 10.1242/dev.197590.

3. Bhattacharya, S., Mukherjee, A., Pisano, S., Dimri, S., Knaane, E., Altshuler, A., Nasser, W., Dey, S., Shi, L., Mizrahi, I., et al. (2023). The biophysical property of the limbal niche maintains stemness through YAP. Cell Death Differ. 30, 1601–1614. 10.1038/s41418-023-01156-7.

4. Gomez, C., Chua, W., Miremadi, A., Quist, S., Headon, D.J., and Watt, F.M. (2013). The Interfollicular Epidermis of Adult Mouse Tail Comprises Two Distinct Cell Lineages that Are Differentially Regulated by Wnt, Edaradd, and Lrig1. Stem Cell Rep. 1, 19–27. 10.1016/j.stemcr.2013.04.001.

5. Sada, A., Jacob, F., Leung, E., Wang, S., White, B.S., Shalloway, D., and Tumbar, T. (2016). Defining the cellular lineage hierarchy in the interfollicular epidermis of adult skin. Nat. Cell Biol. 18, 619–631. 10.1038/ncb3359.

6. Wang, S., Drummond, M.L., Guerrero-Juarez, C.F., Tarapore, E., MacLean, A.L., Stabell, A.R., Wu, S.C., Gutierrez, G., That, B.T., Benavente, C.A., et al. (2020). Single cell transcriptomics of human epidermis identifies basal stem cell transition states. Nat. Commun. 11, 4239. 10.1038/s41467-020-18075-7.

7. Jones, K.B., Furukawa, S., Marangoni, P., Ma, H., Pinkard, H., D’Urso, R., Zilionis, R., Klein, A.M., and Klein, O.D. (2019). Quantitative Clonal Analysis and Single-Cell Transcriptomics Reveal Division Kinetics, Hierarchy, and Fate of Oral Epithelial Progenitor Cells. Cell Stem Cell 24, 183–192.e8. 10.1016/j.stem.2018.10.015.

8. Bickenbach, J.R. (1981). Identification and behavior of label-retaining cells in oral mucosa and skin. J. Dent. Res. 60 *Spec No C*, 1611–1620. 10.1177/002203458106000311011.

9. Bickenbach, J.R., and Mackenzie, I.C. (1984). Identification and Localization of Label-Retaining Cells in Hamster Epithelia. J. Invest. Dermatol. 82, 618–622. 10.1111/1523-1747.ep12261460.

10. Willberg, J., Syrjänen, S., and Hormia, M. (2006). Junctional Epithelium in Rats Is Characterized by Slow Cell Proliferation. J. Periodontol. 77, 840–846. 10.1902/jop.2006.050213.

11. Asaka, T., Akiyama, M., Kitagawa, Y., and Shimizu, H. (2009). Higher density of label-retaining cells in gingival epithelium. J. Dermatol. Sci. 55, 132–134. 10.1016/j.jdermsci.2009.03.006.

12. Byrd, K.M., Piehl, N.C., Patel, J.H., Huh, W.J., Sequeira, I., Lough, K.J., Wagner, B.L., Marangoni, P., Watt, F.M., Klein, O.D., et al. (2019). Heterogeneity within Stratified Epithelial Stem Cell Populations Maintains the Oral Mucosa in Response to Physiological Stress. Cell Stem Cell 25, 814–829.e6. 10.1016/j.stem.2019.11.005.

13. Seubert, A.C., Krafft, M., Bopp, S., Helal, M., Bhandare, P., Wolf, E., Alemany, A., Riedel, A., and Kretzschmar, K. (2024). Spatial transcriptomics reveals molecular cues underlying the site specificity of the adult mouse oral mucosa and its stem cell niches. Stem Cell Rep. 19, 1706–1719. 10.1016/j.stemcr.2024.10.007.

14. Viswanathan, P., Guvendiren, M., Chua, W., Telerman, S.B., Liakath-Ali, K., Burdick, J.A., and Watt, F.M. (2016). Mimicking the topography of the epidermal–dermal interface with elastomer substrates. Integr. Biol. 8, 21–29. 10.1039/C5IB00238A.

15. Mobasseri, S.A., Zijl, S., Salameti, V., Walko, G., Stannard, A., Garcia-Manyes, S., and Watt, F.M. (2019). Patterning of human epidermal stem cells on undulating elastomer substrates reflects differences in cell stiffness. Acta Biomater. 87, 256–264. 10.1016/j.actbio.2019.01.063.

16. Shen, Z., Liu, Z., Sun, L., Li, M., Han, L., Wang, J., Wu, X., and Sang, S. (2023). Constructing epidermal rete ridges using a composite hydrogel to enhance multiple signaling pathways for the maintenance of epidermal stem cell niche. Acta Biomater. 169, 273–288. 10.1016/j.actbio.2023.07.037.

17. Tumbar, T., Guasch, G., Greco, V., Blanpain, C., Lowry, W.E., Rendl, M., and Fuchs, E. (2004). Defining the Epithelial Stem Cell Niche in Skin. Science 303, 359–363. 10.1126/science.1092436.

18. Squier, C.A., and Kremer, M.J. (2001). Biology of Oral Mucosa and Esophagus. JNCI Monogr. 2001, 7–15. 10.1093/oxfordjournals.jncimonographs.a003443.

19. Jones, K.B., and Klein, O.D. (2013). Oral epithelial stem cells in tissue maintenance and disease: the first steps in a long journey. Int. J. Oral Sci. 5, 121–129. 10.1038/ijos.2013.46.

20. Schlüter, H., Paquet-Fifield, S., Gangatirkar, P., Li, J., and Kaur, P. (2011). Functional Characterization of Quiescent Keratinocyte Stem Cells and Their Progeny Reveals a Hierarchical Organization in Human Skin Epidermis. Stem Cells 29, 1256–1268. 10.1002/stem.675.

21. Iglesias-Bartolome, R., Uchiyama, A., Molinolo, A.A., Abusleme, L., Brooks, S.R., Callejas-Valera, J.L., Edwards, D., Doci, C., Asselin-Labat, M.-L., Onaitis, M.W., et al. (2018). Transcriptional signature primes human oral mucosa for rapid wound healing. Sci. Transl. Med. 10, eaap8798. 10.1126/scitranslmed.aap8798.

22. Suebsamarn, O., Kamimura, Y., Suzuki, A., Kodama, Y., Mizuno, R., Osawa, Y., Komatsu, T., Sato, T., Haga, K., Kobayashi, R., et al. (2022). In-process monitoring of a tissue-engineered oral mucosa fabricated on a micropatterned collagen scaffold: use of optical coherence tomography for quality control. Heliyon 8, e11468. 10.1016/j.heliyon.2022.e11468.

23. Coolen, N.A., Verkerk, M., Reijnen, L., Vlig, M., Van Den Bogaerdt, A.J., Breetveld, M., Gibbs, S., Middelkoop, E., and Ulrich, M.M.W. (2007). Culture of Keratinocytes for Transplantation without the Need of Feeder Layer Cells. Cell Transplant. 16, 649–661. 10.3727/000000007783465046.

24. Peters, V.A., Joesting, J.J., and Freund, G.G. (2013). IL-1 receptor 2 (IL-1R2) and its role in immune regulation. Brain. Behav. Immun. 32, 1–8. 10.1016/j.bbi.2012.11.006.

25. Morinaga, H., Mohri, Y., Grachtchouk, M., Asakawa, K., Matsumura, H., Oshima, M., Takayama, N., Kato, T., Nishimori, Y., Sorimachi, Y., et al. (2021). Obesity accelerates hair thinning by stem cell-centric converging mechanisms. Nature 595, 266–271. 10.1038/s41586-021-03624-x.

26. Shyer, A.E., Huycke, T.R., Lee, C., Mahadevan, L., and Tabin, C.J. (2015). Bending Gradients: How the Intestinal Stem Cell Gets Its Home. Cell 161, 569–580. 10.1016/j.cell.2015.03.041.

27. Kraiczy, J., McCarthy, N., Malagola, E., Tie, G., Madha, S., Boffelli, D., Wagner, D.E., Wang, T.C., and Shivdasani, R.A. (2023). Graded BMP signaling within intestinal crypt architecture directs self-organization of the Wnt-secreting stem cell niche. Cell Stem Cell 30, 433–449.e8. 10.1016/j.stem.2023.03.004.

28. Sumigray, K.D., Terwilliger, M., and Lechler, T. (2018). Morphogenesis and Compartmentalization of the Intestinal Crypt. Dev. Cell 45, 183–197.e5. 10.1016/j.devcel.2018.03.024.

29. Walko, G., Woodhouse, S., Pisco, A.O., Rognoni, E., Liakath-Ali, K., Lichtenberger, B.M., Mishra, A., Telerman, S.B., Viswanathan, P., Logtenberg, M., et al. (2017). A genome-wide screen identifies YAP/WBP2 interplay conferring growth advantage on human epidermal stem cells. Nat. Commun. 8, 14744. 10.1038/ncomms14744.

30. Shen, Z., Sun, L., Liu, Z., Li, M., Cao, Y., Han, L., Wang, J., Wu, X., and Sang, S. (2023). Rete ridges: Morphogenesis, function, regulation, and reconstruction. Acta Biomater. 155, 19–34. 10.1016/j.actbio.2022.11.031.

31. Roger, M., Fullard, N., Costello, L., Bradbury, S., Markiewicz, E., O’Reilly, S., Darling, N., Ritchie, P., Määttä, A., Karakesisoglou, I., et al. (2019). Bioengineering the microanatomy of human skin. J. Anat. 234, 438–455. 10.1111/joa.12942.

32. Weinmüllner, R., Zbiral, B., Becirovic, A., Stelzer, E.M., Nagelreiter, F., Schosserer, M., Lämmermann, I., Liendl, L., Lang, M., Terlecki-Zaniewicz, L., et al. (2020). Organotypic human skin culture models constructed with senescent fibroblasts show hallmarks of skin aging. Npj Aging Mech. Dis. 6, 4. 10.1038/s41514-020-0042-x.

33. Suzuki, A., Kodama, Y., Miwa, K., Kishimoto, K., Hoshikawa, E., Haga, K., Sato, T., Mizuno, J., and Izumi, K. (2020). Manufacturing micropatterned collagen scaffolds with chemical-crosslinking for development of biomimetic tissue-engineered oral mucosa. Sci. Rep. 10, 22192. 10.1038/s41598-020-79114-3.

34. Suzuki, A., Kato, H., Kawakami, T., Kodama, Y., Shiozawa, M., Kuwae, H., Miwa, K., Hoshikawa, E., Haga, K., Shiomi, A., et al. (2020). Development of microstructured fish scale collagen scaffolds to manufacture a tissue-engineered oral mucosa equivalent. J. Biomater. Sci. Polym. Ed. 31, 578–600. 10.1080/09205063.2019.1706147.

35. Diamond, I., Owolabi, T., Marco, M., Lam, C., and Glick, A. (2000). Conditional Gene Expression in the Epidermis of Transgenic Mice Using the Tetracycline-Regulated Transactivators tTA and rTA Linked to the Keratin 5 Promoter. J. Invest. Dermatol. 115, 788– 794. 10.1046/j.1523-1747.2000.00144.x.

36. Taniguchi, H., He, M., Wu, P., Kim, S., Paik, R., Sugino, K., Kvitsani, D., Fu, Y., Lu, J., Lin, Y., et al. (2011). A Resource of Cre Driver Lines for Genetic Targeting of GABAergic Neurons in Cerebral Cortex. Neuron 71, 995–1013. 10.1016/j.neuron.2011.07.026.

